# Kinetics and Mechanism of Fentanyl Dissociation from the *µ*-Opioid Receptor

**DOI:** 10.1101/2021.04.25.441335

**Authors:** Paween Mahinthichaichan, Quynh N. Vo, Christopher R. Ellis, Jana Shen

## Abstract

Driven by illicit fentanyl, opioid related deaths have reached the highest level in 2020. Currently, an opioid overdose is resuscitated by the use of naloxone, which competitively binds and antagonizes the *µ*-opioid receptor (mOR). Thus, knowledge of the residence times of opioids at mOR and the unbinding mechanisms is valuable for assessing the effectiveness of naloxone. In the present study we calculate the fentanyl-mOR dissociation time and elucidate the mechanism by applying an enhanced sampling molecular dynamics (MD) technique. Two sets of metadynamics simulations with different initial structures were performed while accounting for the protonation state of the conserved H297^6.52^, which has been suggested to modulate the ligand-mOR affinity and binding mode. Surprisingly, with the N*δ*-protonated H297^6.52^, fentanyl can descend as much as 10 Å below the level of the conserved D147^3.32^ before escaping the receptor, and has a calculated residence time *τ* of 38 s. In contrast, with the N - and doubly protonated H297^6.52^, the calculated *τ* are 2.6 s and 0.9 s, respectively. Analysis suggests that formation of the piperidine–Hid297 hydrogen bond strengthens the hydrophobic contacts with the transmembrane helix (TM) 6, allowing fentanyl to explore a deep pocket. Considering the experimental *τ* of ∼4 min for fentanyl and the role of TM6 in mOR activation, we suggest that the deep insertion mechanism may be biologically relevant. The work paves the way for large-scale computational predictions of opioid dissociation rates to inform the regulatory decision. The profound role of the histidine protonation state found here may shift the paradigm in computational studies of ligand-receptor kinetics.

## INTRODUCTION

Opioids are powerful painkillers but also among the most abused drugs.^1,2^ In 2017, the US Department of Health and Human Services declared the opioid crisis a public health emergency. According to data from the Centers for Disease Control and Prevention,^3^ a record number of 81,000 drug overdose deaths occurred in 2019-2020 and the main driver of the increase is synthetic opioids, primarily illicitly manufactured fentanyl, which is often pressed into counterfeit pills or mixed with heroin or cocaine. Fentanyl is a controlled prescription narcotic and depending on the route of administration 10-400 times more potent than morphine.^4^ The significant rise in overdose deaths from illicitly produced nonpharmaceutical fentanyl has been attributed to fentanyl’s high potency, fast onset, and low manufacturing cost.^4,5^ Additionally, the ease of structural modifications has led to the emergence of illicit fentanyl analogs,^5^ which may have even higher potency and abuse potential.

The main molecular target of fentanyl is the *µ*-opioid receptor (mOR),^6^ a class A G-protein coupled receptor (GPCR) comprised of seven transmembrane (TM) helices (Fig. 1a and b). Fentanyl binds mOR, which induces conformational changes and enabling the receptor to associate with G-protein and *β*-arrestin and activate the related signaling pathways. Current treatment of opioid overdose relies on the use of naloxone, an antagonist of mOR. At the microscopic level, naloxone resuscitates an opioid overdose by competitive binding and deactivation of mOR. Thus, the residence time of an opioid at mOR is an important parameter for evaluating the effectiveness of naloxone or other counter-measures. Measurement of opioid dissociation rates is very costly and non-trivial, as it involves radioligand labeling. Developing a computational capability that can accurately predict residence times and unbinding mechanisms is of significant value.

**Figure 1.**
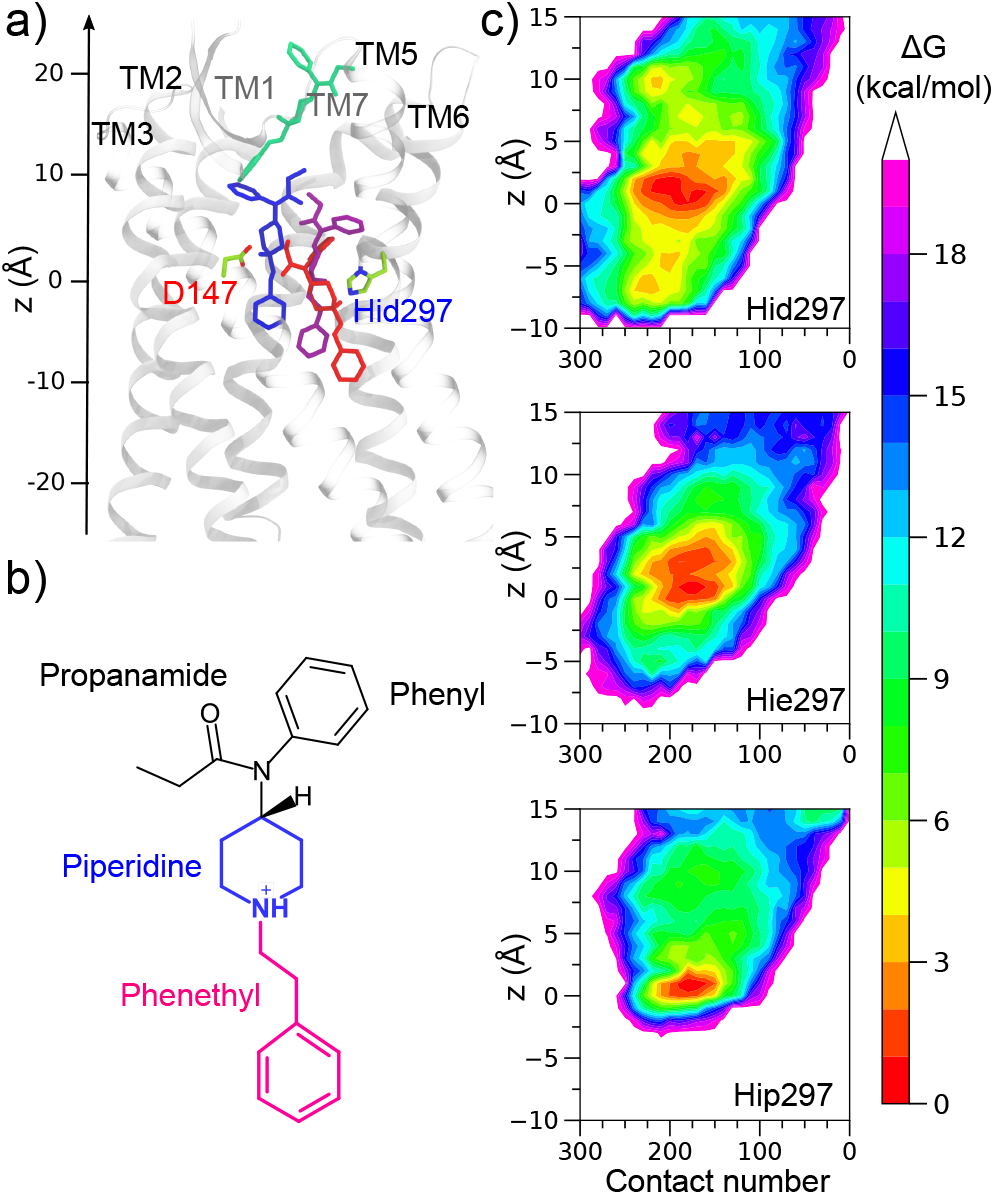
Fentanyl unbinding depends on the protonation state of H297^6.52^. **a)** Snapshots of fentanyl from a metadynamics trajectory with Hid297. Fentanyl is colored according to the time: blue (starting), purple, red (deep pocket), and green (out in solution). The z axis is shown with the origin placed at the C*α* atom of D147^3.32^. **b**) Chemical structure of fentanyl. **c**) Unbiased approximate FE surfaces as a function of fentanyl’s contact number with mOR and its COM z position in the presence of Hid297 (top), Hie297 (middle), and Hip297 (bottom). The trajectories initiated from the global minimum state were used.

An experimental structure of mOR bound to fentanyl or fentanyl-like opioid is lacking; however, the X-ray co-crystal structures of mOR in complex with the morphinan agonist BU72^7^ and antagonist *β*-FNA,^8^ as well as the cryogenic electron microscopy (cryo-EM) model of mOR in complex with an endogenous peptide analog DAMGO^9^ were recently obtained. These structures revealed an orthosteric binding site common to class A GPCRs,^10^ which is located near the center of the receptor, involving transmembrane helices (TMs) 2, 3, 5, 6 and 7 (Fig. 1a). In this site, the ligand forms a salt bridge between the piperidine amine and D147^3.32^ on TM3 (superscript refers to the Ballesteros-Weinstein numbering^11^), consistent with an earlier mutagenesis experiment demonstrating that alanine substitution of D147^3.32^ reduces mOR’s affinity for morphine, DAMGO, and naloxone.^12^ Fentanyl also possesses a piperidine in its core (Fig. 1b), and the piperidine-D147^3.32^ binding mode has been recently validated by molecular docking^13^ and molecular dynamics (MD) simulations.^14–16^ Nonetheless, fentanyl and morphine differ significantly in their structures and signaling biases, which has been speculated as a result of different binding and activation mechanisms.^5^

Using multiple MD techniques including the continuous constant pH and weighted-ensemble (WE) methods,^16^ we recently found that fentanyl can adopt a secondary binding mode through hydrogen bonding between the piperidine and the conserved H297^6.52^ on TM6. Intriguingly, the H297-binding mode only occurred in the simulations with the N*δ*-protonated state.^16^ The protonation state of H297^6.52^ has been repeatedly suggested as important for mOR’s ligand recognition and activity; however, the precise role of H297^6.52^ remains unclear. Mutagenesis experiments showed that at low pH whereby H297^6.52^ is likely protonated, mOR’s affinities for DAMGO and naloxone are reduced;^12,17^ however, the measurement conducted with fentanyl was inconclusive.^18^

MD simulations have been increasingly applied to the study of ligand-receptor kinetics.^19–23^ In this work we employed an enhanced sampling MD technique called well-tempered metadynamics^24,25^ to examine fentanyl’s dissociation mechanism and kinetics as well as to further probe the role of H297^6.52^. In metadynamics, bias potentials are deposited along chosen collective variables to overcome free energy barriers. As such metadynamics is an efficient method for exploring protein-ligand unbinding kinetics,^26–28^ bridging the microsecond simulation and the second-to-minute experimental timescales. Most recently, metadynamics simulations of morphine and buprenorphine dissociation from mOR showed that the ligand transitioned to the vestibule region after the D147^3.32^ salt bridge was disrupted.^23^

To examine fentanyl’s unbinding kinetics and explore the effect of H297^6.52^ protonation state, here we performed two sets of simulations starting from two fentanyl-bound mOR structures and with different H297^6.52^ protonation states. Both sets of simulations demonstrated that fentanyl can insert deep into mOR in the presence of the N*δ*-protonated H297^6.52^ and the corresponding dissociation time is nearly one or two orders of magnitude longer than that with the N_*ϵ*_-protonated or the doubly protonated H297^6.52^, respectively. This work represents a first step towards large-scale predictive modeling of the dissociation kinetics of fentanyl analogs, which may be used to assist the evaluation of the effectiveness of naloxone for opioid overdose treatments.

## RESULTS AND DISCUSSION

### Fentanyl-mOR dissociation pathways are dependent on the protonation state of H297^6.52^

To study fentanyl un-binding, two sets of well-tempered metadynamics^25^ simulations were performed with the three protonation states of H297^6.52^: Hid (proton on N*δ*), Hie (proton on N_*ϵ*_), and Hip (doubly protonated). In the first set of simulations comprised of 48 independent metadynamics runs for each protonation state, a MD-relaxed docked structure was used, wherein fentanyl forms a salt-bridge with D147^3.32^ and its center-of-mass (COM) z is about 4.5 Å relative to the C*α* z of D147^3.32^, similar to BU72 in the crystal structure (PDB: 5C1M).^7^ The second set of simulations comprised of 15 independent metadynamics runs for each protonation state was initiated from the most populated configuration of the fentanyl-mOR complex from the recent 40-*µ*s weighted-ensemble (WE) simulations.^16^ In this structure, fentanyl also forms a salt-bridge with D147^3.32^ but the COM z is about 1 Å. Note, the C*α* z of D147^3.32^ was stable in the simulations (e.g., standard deviation of 0.3 Å in the Hid trajectories). For brevity, we will refer to this starting structure as the global minimum state. To drive the dissociation of fentanyl, a biasing potential was deposited every 10 ps along two collective variables: fentanyl’s COM z position and the contact number between fentanyl and mOR heavy atoms (defined in Eq. 1). The biased metadynamics time of dissociation ranged from 10 to 90 ns. For conciseness, we will omit the Ballesteros-Weinstein numbering^11^ in the discussion.

We first discuss the simulations started from the global minimum state (Fig. 1a,b, and Fig. S1). In the presence of Hid/Hie297, fentanyl moved below the level of D147 (z=0) before existing the receptor; however, it stayed in the global minimum region in the presence of Hip297. We calculated the unbiased approximate free energy (FE) surfaces projected onto fentanyl’s COM z position (relative to the C*α* atom of D147) and the heavy-atom contact number with mOR for trajectories with the three protonation states of H297 (Fig. 1c). Regardless of the protonation state, the FE surfaces show a similarly located minimum region representing the D147-bound state; however, with Hip297 the minimum region is more restricted and fentanyl remains above -2 Å, suggesting a more tightly bound state. With Hid/Hie297, the FE surfaces display a broader minimum region, suggesting a more flexible D147-bound state, and surprisingly, fentanyl samples regions in mOR with z down to ∼-10 with Hid297 or ∼-7 Å with Hie297. Closer examination of the FE surface with Hid297 reveals that fentanyl’s deep insertion (defined as below -5 Å) is facilitated by a larger number of contacts with mOR as compared to the D147-bound state. Importantly, there are local minima at z values of ∼-5 Å and ∼-7 Å, which do not exist in the FE surface with Hie297. This suggests that the Hid tautomer induces unique interactions that can stabilize fentanyl in a deep pocket (see later discussion).

We now turn to the simulations initiated from the relaxed docked structure where fentanyl’s z is ∼4.5 Å. In the majority of the trajectories (60-80% with the different H297 protonation states), fentanyl directly exited after briefly sampling the weakly bound state and without sampling the global free energy minimum (FE surfaces are given in Fig. S2). This analysis suggests that the “direct exit” trajectories are artifact and should be discarded. Interestingly, the FE surfaces of other trajectories revealed that the global minimum state was sampled (Fig. S2), consistent with the simulations initiated from the global minimum state. Furthermore, in these trajectories, fentanyl sampled the regions with the z values as low as ∼-10 Å in the presence of Hid297 or ∼-8 Å in the presence of Hie297 (Fig. S2), consistent with the other set of simulations.

### Fentanyl-mOR dissociation time is dependent on the protonation state of H297^6.52^

Following the work of others,^23,27,29^ we analyzed the unbiased fentanyl escape times recovered from the individual trajectories using the Poisson distribution (Fig. S3). By fitting to the theoretical cumulative distribution function,^29^ the fentanyl-mOR dissociation time (also called fentanyl residence time) *τ* (*τ* = 1*/k*_off_) was estimated. Based on the simulations started from the global minimum state, the calculated *τ* values are 38±19 s, 2.6±0.9 s, and 0.9±0.2 s in the presence of Hid297, Hie297, and Hip297, respectively (Table 1 and Fig. S3). In good agreement with these values, the trajectories initiated from the relaxed docked structure that visited the global minimum state yielded the respective *τ* values of 27±12 s, 6.1±1 s, and 0.6±0.2 s (Table 1 and Fig. S4).

**Table 1.** Calculated fentanyl-mOR dissociation times with different protonation states of H297^6.52^ from the two sets of simulations

| H297<sup>6,52</sup> | Global minimum | Relaxed docked |
| --- | --- | --- |
| Hid | $38 \pm 19$ s | $27 \pm 12$ s |
| Hie | $2.6 \pm 0.9$ s | $6 \pm 1$ s |
| Hip | $0.9 \pm 0.2$ s | $0.6 \pm 0.2$ s |

Errors are from the boot-strap analysis. Relaxed docked refer to the trajectories started from the relaxed docked structure that sampled the global minimum state.

The calculated fentanyl residence time of 38±19 s in the presence of Hid297 (or 27±12 s based on the simulations started from the docked structure) is within one order of magnitude from the recently measured value of ∼4 min by Li and colleagues from FDA (data published on GitHub https://github.com/FDA/Mechanistic-PK-PD-Model-to-Rescue-Opiod-Overdose).

### Deep insertion is related to fentanyl’s interaction with H297^6.52^

Since our previous work employing the WE and unbiased equilibrium MD simulations suggested that fentanyl can form either a salt bridge with D147 or a hydrogen bond with Hid297,^16^ we analyzed the fentanyl interactions with D147 and H297 using the Hid, Hie, and Hip trajectories initiated from the global minimum state. The approximated FE surfaces are projected onto the FEN–D147 and FEN–H297 distances. We also plotted the volumetric occupancy maps of piperidine (which forms interactions with D147 or H297) and phenethyl group (at the bottom of the fentanyl structure) to gain a visual impression of the location of fentanyl throughout the dissociation process.

The occupancy maps based on the trajectories with Hid297, Hie297 and Hip297 show clear differences. Consistent with the FE surfaces in terms of fentanyl’s z position and fentanyl-mOR contact number (Fig. 1c), fentanyl appears to sample a larger volume and its phenethyl group inserts deeper into mOR in the presence of Hid297 (Fig. 2a,c-d). The occupancy map of the Hip trajectories shows that fentanyl does not reach H297, which may be explained by the repulsion between the positively charged piperidine and Hip297, consistent with the previous equilibrium MD and constant pH MD simulations.^16^

**Figure 2.**
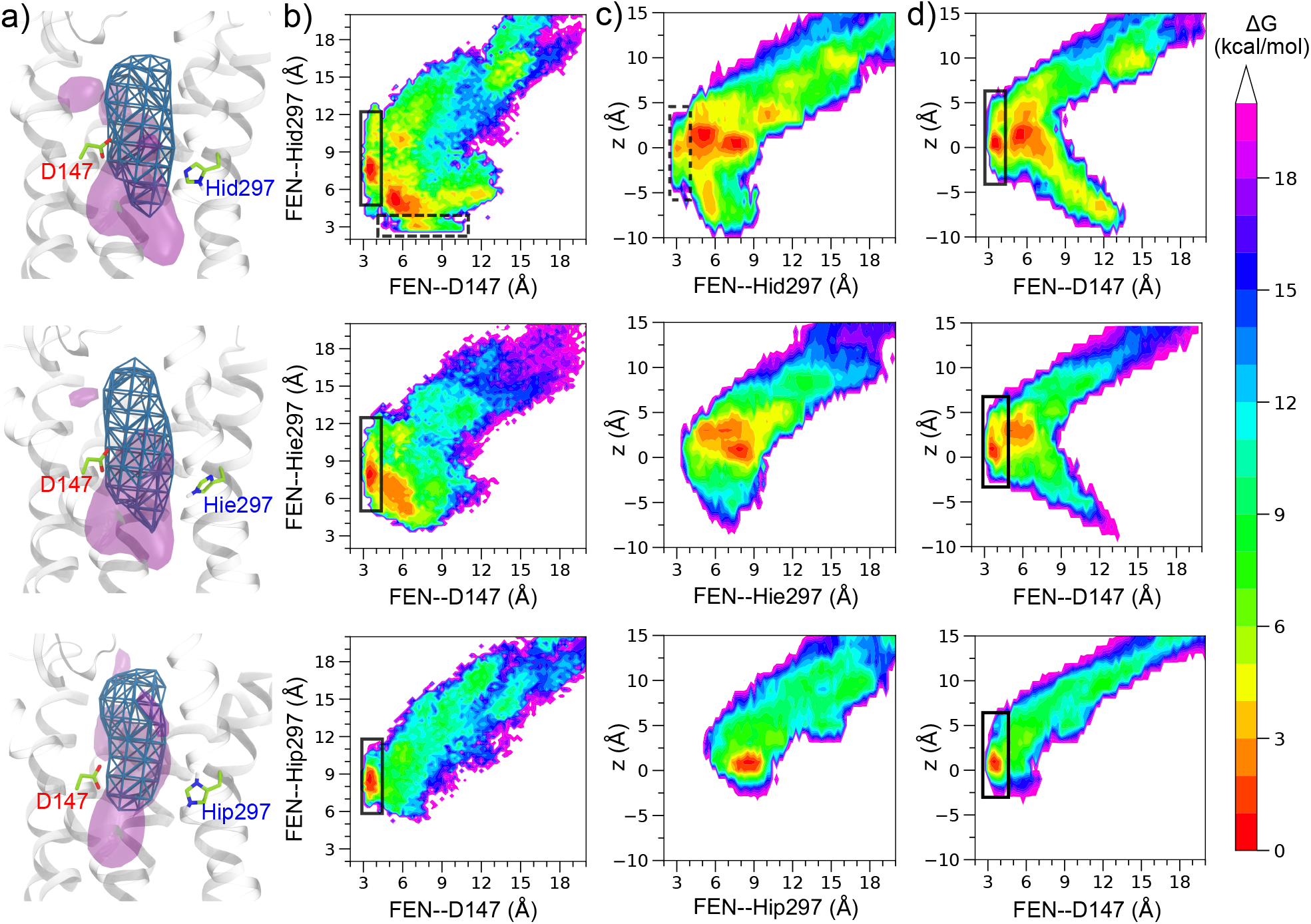
Fentanyl’s location and interactions with D147^3.32^/H297^6.52^ are dependent on the protonation state of H297^6.52^. **a)** Volumetric occupancy map of the piperidine (green mesh) and phenethyl (purple) groups from the Hid (**top**), Hie (**middle**), or Hip (**bottom**) trajectories. Calculations were performed with the VolMap plugin in VMD^30^ (occupancy cutoff of 0.03). The isosurface maps were rendered with VMD.^30^ **b)** Approximate FE surfaces projected onto fentanyl’s distances to D147 and H297 based on the Hid (**top**), Hie (**middle**), and Hip (**bottom**) trajectories. The FEN–D147 distance is measured between piperidine:N and D147:C*γ*. The FEN–H297 distance is measured between piperidine:N and H297:N_*ϵ*_ or N*δ* (the closest). **c**,**d)** Approximate FE surfaces projected onto fentanyl’s COM *z* position and distance to H297 (**c**) or D147 (**d**) based on the Hid (**top**), Hie (**middle**), and Hip (**bottom**) trajectories. The solid and dashed black boxes highlight the regions where the D147 salt bridge and the H297 hydrogen bond are formed. The FES surfaces were calculated as a Boltzmann average of the free energies of the reweighted individual trajectories. The reweighting protocol 31 in PLUMED 32 was used.

Projected onto the FEN–D147 and FEN–H297 distances, the FE surface based on the Hid trajectories is most diffuse and contains many local minima, whereas the FE surface from the Hip trajectories is most localized and displays only one minimum located around the FEN–D147 distance of 3–4 Å and the FEN–H297 distance of 7–8 Å. This minimum corresponds to the D147-bound state in which the piperidine forms a salt bridge with D147 while the interaction with H297 is negligible. The FE surfaces from the Hid and Hie trajectories show an additional minimum region with somewhat increased FEN–D147 distance (4–6 Å) and decreased FEN–H297 distance (4.5–6 Å), suggesting that fentanyl can form van der Waals interactions with both D147 and H297 in the presence of Hid297 or Hie297. A major difference between the two FES surfaces is that the former contains a local minimum at the FEN–D147 distance of 7 Å and the FEN–H297 distance of 3 Å, while the latter does not extend below the FEN–H297 distance of 4 Å. This is in agreement with our previous WE simulations^16^ which showed that fentanyl can form a hydrogen bond with Hid297 but not Hie297 (Fig. 2b, dashed black box regions).

The FE surfaces in terms of the FEN–D147 or FEN–H297 distance and fentanyl’s z position show that when the FEN– D147 salt bridge is formed, fentanyl’s z varies from -2 to 5 Å (Fig. 2d and S6a); in contrast, when the FEN–Hid297 hydrogen bond is formed, fentanyl’s position is lower and z varies from -5 to 3 Å (Fig. 2c top and S6b left). This is readily understood given the lower position of the imidazole ring relative to the carboxylate.^16^ Curiously, the FE surface of the Hid trajectories also displays a local minimum at the FEN–Hid297 distance of 5 to 6 Å and z value of -8 to -4 Å (Fig. 2c top and S6b left), which indicates that in the deeply inserted state, fentanyl is not hydrogen bonded with Hid297.

### Fentanyl-TM6 interactions are weakened with Hip297

To further understand how the H297 protonation state impacts fentanyl’s sampling in mOR, we calculated the occupancies of contacts between the fentanyl constituent groups and mOR residues based on the Hid, Hie, and Hip trajectories initiated from the global minimum state (Fig. 3). The contact profiles with Hid297 and Hie297 are similar; whereas the profile with Hip297 contains a significantly lower number of contacts. Most noticeably, the piperidine–TM6 interactions are nearly abolished (occupancy below 10%), which may be explained by the electrostatic repulsion between the positively charged piperidine and imidazole rings. By contrast, with Hid/Hie297 the piperidine ring interacts with several TM6 residues including V300, I296, H297, and W293, which have the descending C*α* z positions of approximately 7, 2.5, 2, - 3.5 Å, respectively. Note, although the C*α* z of H297 is ∼2 Å higher than D147, its sidechain z position is lower. It is worth noticing that the piperidine only briefly interacts with Hie297 (occupancy below 10%), due to the lack of the hydrogen bond formation as with Hid297 (Fig. 2). In the presence of Hip297, the phenylpropanamide and phenethyl groups also have less or significantly weakened contacts with TM6, e.g., the contact between phenylpropanamide and W293 (C*α* z ∼ -3.5 Å) and between the phenylethyl and F289 (C*α* z -10 Å) are absent. This is related to the fact that fentanyl does not sample the deep pocket with Hip297. Consistently, the piperidine does not interact with M151 (z -4 Å) on TM3. All the aforementioned differences between the Hip and Hid/Hie trajectories are in agreement with the data from the simulations starting from the relaxed docked structure (Fig. S7).

**Figure 3.**
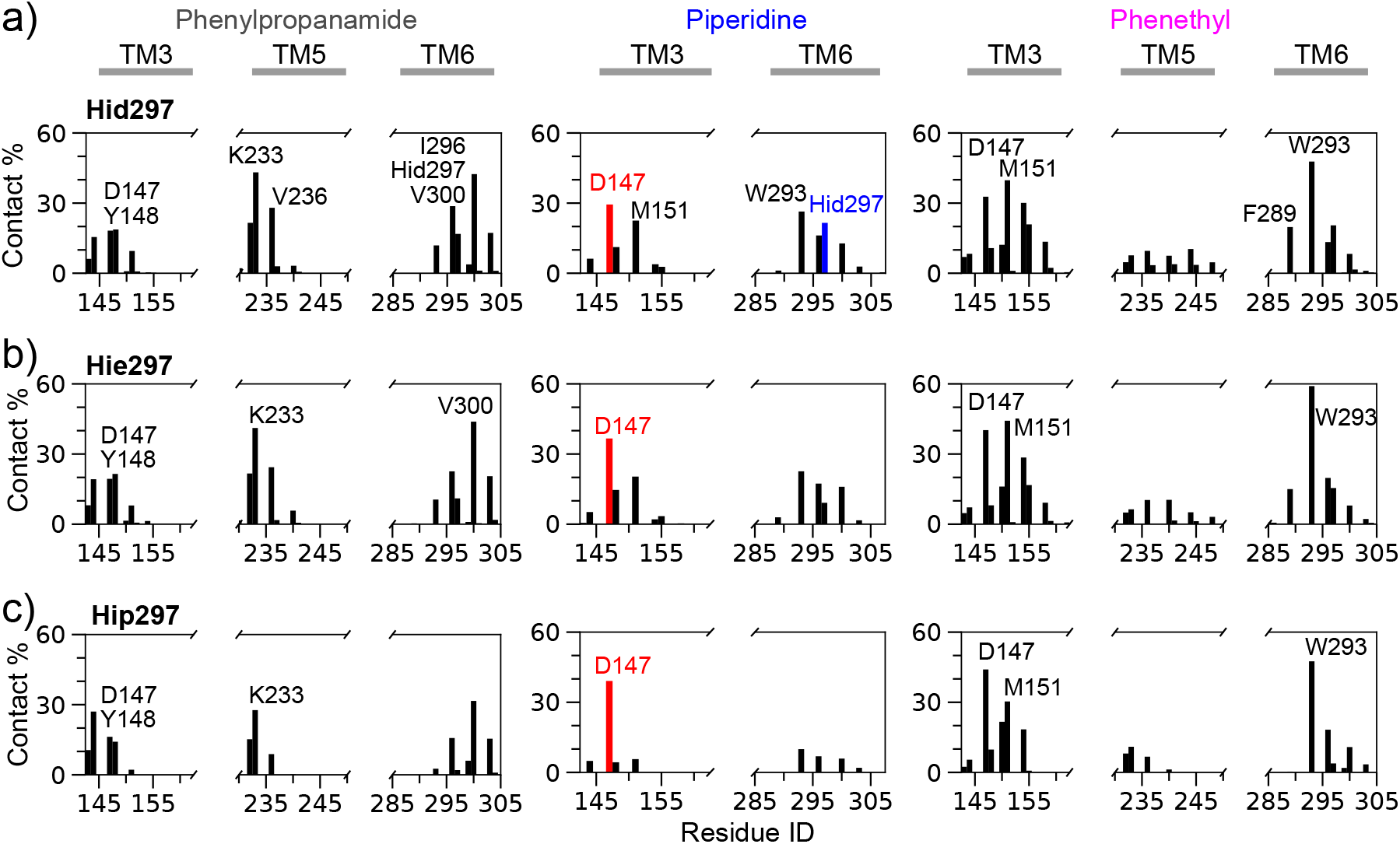
Occupancies of contacts between fentanyl and mOR residues. **a,b,c)** Fraction of contacts between mOR residues and the phenylpropanamide (left), piperidine (middle), or phenethyl (right) group calculated based on the Hid (**a**), Hie (**b**) or Hip (**c**) trajectories initiated from the global minimum state. Residues with fractions ≥20% are labeled. A contact was considered formed if the heavy-atom distance is below 4.5 Å.

### Fentanyl interacts with the deep pocket residues in the presence of Hid297

To further characterize fentanyl’s deep insertion, we calculated the fentanyl-mOR contact profile based on the Hid trajectories in which fentanyl moved below -5 Å (Fig. 4) and compare it against the profiles obtained with all Hid (Fig. 3a) and Hie trajectories (Fig. 3b). For the piperidine group, the most prominent interactions (over 80% occupancy) that are enriched in the deep pocket trajectories are with W293 (z ∼-3.5 Å) on TM6 and M151 (z ∼-4 Å) on TM3. Importantly, the occupancy for the piperidine-H297 contact is over 50% in the deep pocket trajectories, compared to 25 % with all Hid trajectories (Fig. 3a) or under 10% with the Hie trajectories (Fig. 3b). For the phenethyl group, the most prominent interactions are with F289 (z ∼-10 Å) on TM6, S154/I155 (z ∼-9.5 Å) and L158 (z ∼-14 Å) on TM3, as well as P244 (z ∼-8 Å) on TM5. Note, I155 and F289 belong to the conserved core triad.^7^ These data suggest that the piperidine-Hid297 hydrogen bond formation strengthens the interaction between piperidine and W293, allowing the phenethyl group to reach deeper into mOR and form hydrophobic contacts with the deep pocket residues located below -5 Å. A related question is whether the deep pocket is dynamically formed in the presence of fentanyl. To address this, we calculated the solvent accessible surface areas of the deep-pocket residues (I155, L158, P244, and F289) from the apo mOR simulations (500 ns equilibrium MD data taken from Ref.^16^) and the holo trajectories with fentanyl’s z position below -5 Å (Table S1). The solvent exposure of the deep-pocket residues, particularly F289, is indeed increased, which suggests that the deep pocket is dynamically formed in the presence of fentanyl.

**Figure 4.**
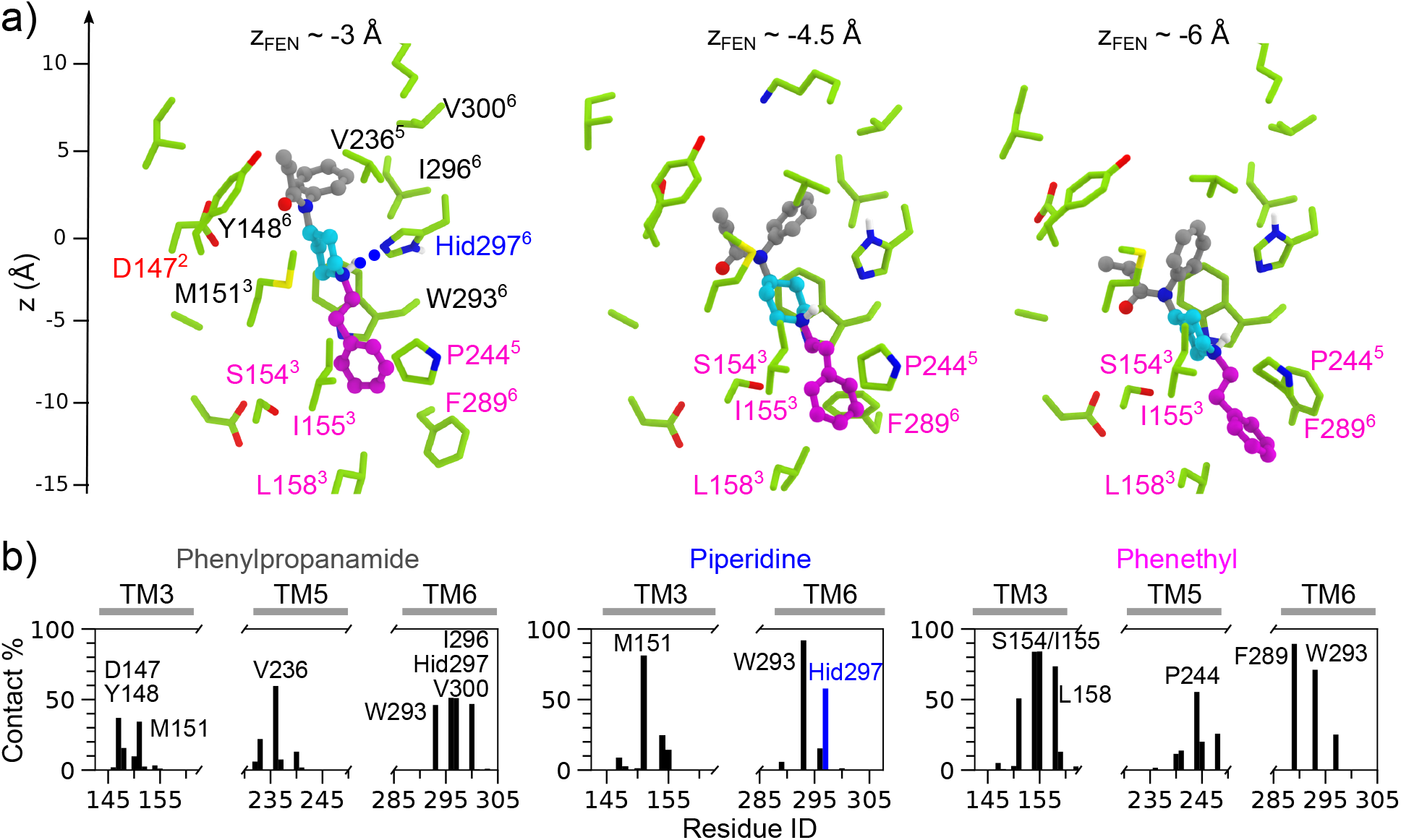
Fentanyl-mOR interactions in the deep pocket. **a)** Snapshots taken from the Hid trajectories initiated from the global minimum state. Left, fentanyl’s z∼-3 Å; the piperidine forms a hydrogen bond with Hid297. Middle, fentanyl’s z∼-4.5 Å; the piperidine forms van der Waals contacts with Hid297. Right, fentanyl’s z∼-6 Å; mainly hydrophobic contacts are formed. The phenylpropanamide, piperidine, and phenethyl groups are colored grey, cyan, and magenta, respectively. Residues contacting fentanyl are shown in the stick model and labeled. Those with a C*α* z position below -5 Å are colored magenta and referred to as the deep pocket residues in the main text. Superscripts refer to the TM helix numbers. In the descending order, the C*α* z positions of V300, I296, H297, W293, and F289 on TM6 are approximately 7, 2.5, 2, -3.5, and -10 Å, respectively, and those of M151, I155, S154, and L158 on TM3 are approximately -4, -9, -9.5 and -14 Å, respectively. The C*α* z position of P244 on TM5 is -8 Å. **b)** Occupancies of the contacts between the phenylpropanamide, piperidine, or phenethyl groups and the mOR residues based on the Hid trajectories initiated from the global minimum state. Residues with contact occupancies ≥30% are labeled. The contacts are defined using a 4.5-Å heavy-atom distance cutoff. Only the trajectories in which fentanyl’s z position went below -5 Å were used.

## CONCLUDING DISCUSSION

In summary, two sets of metadynamics simulations (a total of 189 trajectories) were carried out to investigate the kinetics and mechanism of fentanyl-mOR dissociation and the impact of the H297 protonation state. In the presence of Hid297, we found that fentanyl can descend as much as 10 Å below the C*α* z position of D147 before escaping from the receptor. By contrast, the insertion is shallower (z as low as -7 Å) with Hie297, and it is absent with Hip297. The calculated residence time of fentanyl from the simulations initiated from the global minimum state is 38±19 s, 2.6±0.9 s, or 0.9±0.2 s with Hid297, Hie297, or Hip297, respectively, similar to the simulations initiated from a relaxed docked structure in which fentanyl sampled the global minimum region. Interestingly, the residence time with Hid297 is within one order of magnitude from the recent experimental value of ∼4 min (https://github.com/FDA/Mechanistic-PK-PD-Model-to-Rescue-Opiod-Overdose) and the previously reported value of ∼5 min for carfentanil,^33^ a close analog of fentanyl. Note, the observed time range for ligand-mOR dissociation was reported as 0.1–10 min,^34^ and the residence times of the morphinan compounds buprenorphine and naloxone are about 10 min^35^ and 1 min,^36^ respectively. Considering the agreement with the experimental timescale and the consistency between the two sets of simulations, we suggest that the deep insertion mechanism may be biologically relevant. Furthermore, the calculated fentanyl-mOR dissociation time is similar to the measured residence time of 29 s for the G-protein mimetic Nb39 binding to fentanyl-bound mOR.^37^ Since the full activation of mOR requires coupling of the agonist-bound mOR to a G-protein,^38^ this agreement lends further support to the potential biological relevance of the deep insertion mechanism.

The analysis suggests that fentanyl’s deep insertion (z below -5 Å) is due to the formation of a hydrogen bond between piperidine’s amine nitrogen and N of H297, corroborating our recent WE simulations of fentanyl-mOR binding.^16^ The contact calculations suggest that the fentanyl-H297 hydrogen bond formation strengthens the interaction between the piperidine ring and W293, which is positioned below H297 on TM6 with z ∼-3.5 Å. The data further demonstrates that these interactions stabilize the hydrophobic contacts between fentanyl’s phenethyl group and the deep pocket residues with z positions below -5 Å, including F289 on TM6 (z ∼-10 Å), I155/S154 (z ∼-9/-9.5Å), and L158 (z ∼-14 Å) on TM3. Despite having similar fentanyl-mOR interactions, the lack of hydrogen bonding with H297 weakens the piperidine-TM6 contacts, which results in a shallower insertion of fentanyl and one order of magnitude decrease in the dissociation time with Hie297. In stark contrast, the electrostatic repulsion between the charged piperidine and Hip297 prohibits many interactions with TM6 residues, resulting in a higher z position of fentanyl and a further decrease in the dissociation time. We note, all residues that are involved in the deep pocket contacts are conserved. Given the central role of TM6 in mOR activation, the hydrophobic interactions with TM6 residues such as W293 and F289 may be functionally relevant. Future studies aimed at the conformational transitions of mOR may provide further insights.

The dissociation mechanism of fentanyl uncovered by our study differs significantly from that of morphine and buprenorphine, which were found to dissociate directly from the orthosteric site region in a recent metadynamics study by the Filizola group (15 trajectories, the protonation state of H297^6.52^ is unclear).^23^ This is however not surprising, as fentanyl differs from morphine in many aspects. The elongated shape of fentanyl may allow it to insert deep into mOR, which may not be possible with bulkier ligands such as morphine and analogs. Fentanyl’s deep insertion is consistent with a previous study, which found that acetylcholine (a small elongated molecule) can diffuse into a deep pocket of M3 and M4 muscarinic acetylcholine receptors.^20^ Preclinical pharmacology studies demonstrated that fentanyl displays a bias for *β*-arrestin relative to G-protein signaling, which is in contrast to morphine that displays a modest bias for G protein signaling.^4,39^ The significant pharmacological difference has been speculated as a result of the different mechanism of binding and activation.^5^ The present metadynamics data adds to the evidence in support of this hypothesis.

The present study has several caveats. A potential change in the protonation state of H297 during fentanyl dissociation is neglected. No significant conformational changes of mOR was observed upon fentanyl dissociation, which is expected due to the short simulation timescale. Due to the use of biasing potential and short timescale of metadynamics, the kinetics calculations, unbiased free energies, as well as the contact calculations may be of limited accuracy. Nonetheless, these caveats are not expected to alter our conclusions regarding the mechanism of deep pocket insertion and how the protonation state of H297^6.52^ alters the unbinding kinetic rate. To test the generality of the deep pocket mechanism and to further explore the role of H297^6.52^, a systematic study of fentanyl and morphine analogs is underway. In addition to the mechanistic insights, our work provides a computational protocol that can be used to predict the dissociation rates of opioids to assist the regulatory evaluation of strategies for drug overdose reversal. The profound role of histidine protonation state discovered here may shift the paradigm in computational studies of ligand-receptor kinetics which is gaining increased attention.

### Methods and Protocols

The protein and lipids (POPC (1-palmitoyl-2-oleoyl-glycero-3-phosphocholine) and cholesterol) were represented by the CHARMM c36m protein^40,41^ and c36 lipid^42^ force fields, respectively. Water was represented by the CHARMM-style TIP3P force field.^43^ Fentanyl was represented by the CGenFF force field (version 3.0.1) obtained through the ParamChem server.^44^ Using NAMD 2.13,^45^ we first carried out 100-ns MD simulations of the membrane-embedded apo mOR based on the BU72-mOR complex crystal structure (PDB:5C1M)^7^ at 1 atm and 310 K. The protonation states were determined by the membrane-enabled hybrid-solvent continuous constant pH molecular dynamics (CpHMD) method with pH replica exchange^46,47^ in CHARMM.^43^ A fentanyl-bound mOR model was prepared by superimposing a top fentanyl binding pose obtained from a previous docking study^13^ onto the equilibrated structure of membrane-embedded apo mOR. The resulting model of the fentanyl-mOR complex was then relaxed by 65-ns NPT simulations at 1 atm and 310 K and used as the starting structure for the first set of simulations comprised of 48 independent metadynamics runs for each H297^6.52^ protonation state. The second set of simulations comprised of 15 metadynamics runs for each H297^6.52^ protonation state was initiated from the most probable configuration (global minimum state) of the fentanyl-mOR complex taken from our recent WE trajectories.^16^ Together, a total of 189 simulations with an aggregate time of ∼6 *µ*s were performed.

The well-tempered metadynamics simulations^25^ were performed for the membrane-embedded fentanyl-mOR complex using the Collective Variables (ColVars) module^48^ in NAMD 2.13.^45^ The COM z position of fentanyl relative to that of the orthosteric site C*α* atoms (defined in Supporting Information) and the fentanyl-mOR contact number (CN) were used as the collective variables.

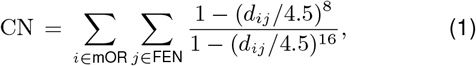

where *d*_*ij*_ is the distance between the heavy atoms *i* and *j* in mOR and fentanyl, respectively. The effective distance cutoff is 4.5 Å. A Gaussian bias potential was deposited every 10 ps, following the previous metadynamics studies of proteinligand unbinding kinetics.^27^ Note, our test simulations with 20 or 30 ps deposition time did not result in significant differences in the calculated dissociation times.

## Supporting Information Available

Detailed methods, protocols, and additional figures are provided in Supporting Information.

## Acknowledgement

P.M. and Q.N.V. are supported by ORISE fellowships through an interagency agreement between the Department of Energy and FDA. Financial support from the National Institutes of Health (R01GM098818) to J.S. is acknowledged. We thank Pratyush Tiwary (University of Maryland, College Park) for the helpful discussion of metadynamics simulations. We thank Lidiya Stavitskaya (FDA) for the discussion of the opioids and experimental data. We thank David G. Strauss, Zhihua Li and John Mann (FDA) for sharing their kinetic measurement data of fentanyl. Disclaimer: the findings and conclusions in this manuscript have not been formally disseminated by the FDA and should not be construed to represent any agency determination or policy.

## Graphical TOC Entry

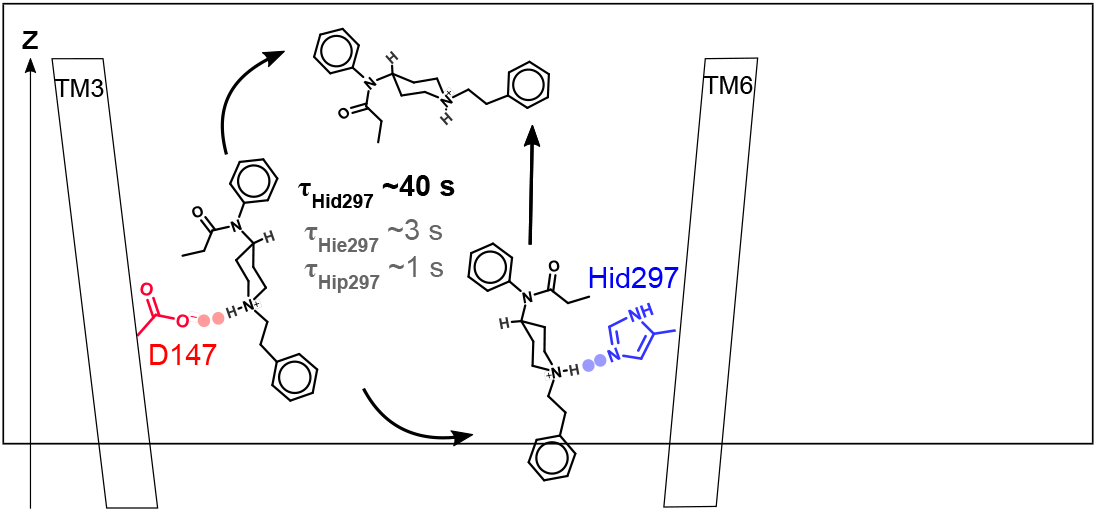

