## Supplemental for "Kinetics and Mechanism of Fentanyl Dissociation from the *µ*-Opioid Receptor"

### Contents

### List of Figures


S1 Starting structures for the two sets of simulations . . . . . . . . . . . . . . . . S12

S2 Free energy surfaces as a function of fentanyl's contact number and z position based on the first set of simulations . . . . . . . . . . . . . . . . . . . . . S13

S3 Poisson fitting analysis of the fentanyl-mOR dissociation times based on the first set of simulations . . . . . . . . . . . . . . . . . . . . . . . . . . . . . S14

S4 Poisson fitting analysis of the fentanyl-mOR dissociation times based on the second set of simulations . . . . . . . . . . . . . . . . . . . . . . . . . . . S15

S5 Robustness of the cutoff used for defining the unbound state . . . . . . . . S16

S6 Free energies as a function of fentanyl's z and distance to D147 or H297 from the relaxed docked structure . . . . . . . . . . . . . . . . . . . . . . . . . . S17

S7 Fentanyl-mOR contact profile from the second set of simulations . . . . . . S18

### Methods and Protocols

**Summary.** The protein and lipids (POPC (1-palmitoyl-2-oleoyl-glycero-3-phosphocholine) and cholesterol) were represented by the CHARMM c36m and c36 force fields,[S1–S3] respectively. Water was represented by the CHARMM-style TIP3P force field.[S4] Fentanyl was represented by the CGenFF force field (version 3.0.1) obtained through the ParamChem server.[S5]

Using NAMD 2.13,[S6] We first carried out 100-ns MD simulations of the the membrane-embedded apo mOR based on the BU72-mOR complex crystal structure (PDB:5C1M)[S7] at 1 atm and 310 K. The protonation states were determined by the membrane-enabled hybrid-solvent continuous constant pH molecular dynamics (CpHMD) method with pH replica exchange[S8,S9] in CHARMM.[S4] A fentanyl-bound mOR model was prepared by superimposing a top fentanyl binding pose obtained from a previous docking study[S10] onto the equilibrated structure of membrane-embedded apo mOR. The resulting model of the fentanyl-mOR complex was then relaxed in 65-ns NPT simulations at 1 atm and 310 K and used as a starting structure for the equilibration simulation before metadynamics.

The well-tempered metadynamics simulations[S11] were performed for the membrane-embedded fentanyl-mOR complex using the Collective Variables (ColVars) module[S12] in NAMD 2.13.[S6] 48 independent metadynamics runs were performed with each of the 3 protonation states of H297$^{6.52}$ (Hid, Hie, and Hip), amounting to a total of 144 simulations with an aggregate time of ~4 $\mu$s. The center-of-mass $z$ position of fentanyl relative to that of the orthosteric site C$\alpha$ atoms (see Fig. S1) and the fentanyl-mOR contact number (CN, defined in Eq.1) were used as the collective variables. A Gaussian bias potential was deposited every 10 ps, as in the previous metadynamics studies of inhibitor-kinase unbinding kinetics.[S13,S14] Note, our test simulations with 20 or 30 ps deposition time did not result in significant differences in the calculated dissociation times.

**Preparation and equilibration of the membrane-embedded apo active mOR structure.** The X-ray crystal structure of mouse $\mu$-opioid receptor (mOR) in complex with a morphinan agonist BU72 and G-protein memetic Nb39 (PDB ID: 5C1M)[S7] was used as the starting model for the apo mOR simulation. BU72 and Nb39 were removed, while the X-ray resolved cholesterol molecule bound at an extracellular interface between transmembrane helix 6 (TM6) and TM7 and all X-ray resolved water molecules were kept. A cysteine-s-acetamide at position 57 was converted to a cysteine (Cys57). Seven water molecules were added to the protein using the DOWSER program with the default threshold energy of -10 kcal/mol.[S15] The apo mOR structure was oriented with respect to the membrane using the Orientations of Proteins in Membranes database.[S16] A disulfide bond was imposed between residues C140 and C217. All titratable residues were fixed in the standard protonation states (charged Asp, Glu, and neutral His), and histidines were in the Hid neutral tautomer form, as in the default setting of CHARMM-GUI[S17] (see later). The CHARMM-GUI web server[S17] was then used to assemble the system of mOR embedded in a solvated POPC (1-palmitoyl-2-oleoyl-glycero-3-phosphocholine) lipid bilayer. The membrane-embedded apo mOR comprised ~79,000 atoms, 210 lipid molecules, ~16,000 water molecules, and 150 mM NaCl (43 $Na^+$ and 56 $Cl^-$ ions). The initial dimension of the bilayer was $\sim 90\times 90\times 106$ Å$^3$.

The system was energy minimized for 5,000 steps using the conjugate gradient algorithm followed by a series of equilibration simulations for a total of 65 ns with the first 0.2 ns under the NVT and the rest under the NPT ensemble. In the first 0.2 ns, the positions of the protein backbone and sidechain heavy atoms were harmonically restrained with the force constant of 10 and then reduced to 5 and 2.5 kcal/mol/Å$^2$. The oxygen atoms of the bound (X-ray and DOWSER added) water molecules were restrained with the force constant of 5 and then reduced to 2.5 kcal/mol/Å$^2$. The lipid head groups were restrained with the force constant of 5 kcal/mol/Å$^2$. In the next 2.3 ns, the force constant for the protein backbone and sidechain were reduced from 2.5 to 1.25 kcal/mol/Å$^2$. The bound water molecules were restrained with the force constant of 1.25 kcal/mol/Å$^2$. The lipid head groups were restrained with the force constant gradually reduced from 1 to 0.1 kcal/mol/Å$^2$. In the next 2.5 ns, the restraints on the lipids were removed, allowing the lipid bilayer to relax, while those on the bound water molecules and the protein remained. In the next 60 ns, the lipid bilayer continued to relax, with the restraints on the protein removed and those on the bound water molecules remaining. The system achieved the dimension of $\sim$87$\times$87$\times$102 Å$^3$.

All of the above simulations were performed using the NAMD 2.13[S6] molecular dynamics package. The protein and lipids were represented by CHARMM c36m protein and c36 lipid force fields, respectively.[S1–S3] Water was represented by the CHARMM style TIP3P model.[S4] The periodic boundary conditions were used. All bond and angles involving hydrogen atoms were constrained using the SHAKE algorithm[S18] to allow a 2-fs time step. The van der Waals interactions were smoothly switched to zero between 10 and 12 Å. The particle mesh Ewald (PME) method[S19] was used to calculate long-range electrostatic forces with a sixth-order interpolation and a grid spacing of $\sim$1 Å. The temperature was maintained at 310 K by Langevin dynamics with a damping coefficient $\gamma$ of 1 ps$^{-1}$. The simulations under an NPT ensemble were performed in a flexible cell with constant xy ratio, and the modified Nosé-Hoover Langevin piston method with the 200-fs period was used to maintain the pressure at 1 atm.[S20,S21]

**Determination of mOR protonation states by continuous constant pH molecular dynamics (CpHMD).** To determine the protonation states of Asp, Glu, and His sidechains, we performed the pH replica exchange membrane-embedded hybrid-solvent CpHMD simulations[S8,S9,S22] using the CHARMM package.[S4] The detailed protocols can be found here.[S23] The replica-exchange CpHMD simulations were performed, starting from the above equilibrated apo active mOR structure. The pH replica-exchange protocol[S8] included 16 replicas in the pH range 2.5–9.5 with an increment of 0.5 unit. A membrane generalized-Born (GB) calculation[S24] was invoked every 10 MD steps to obtain the solvation forces on the protein before updating the titration coordinates. In the membrane GB calculation, the default settings were used.[S8] Each pH replica underwent MD in the NPT ensemble, with an aggregate sampling time of 320 ns. All Asp, Glu, and His sidechains were allowed to titrate, and the corresponding model p$K_\mathrm{a}$'s are 3.8, 4.2, and 6.5.[S25] Based on the CpHMD titration, we set all Asp and Glu sidechains charged (including D114 which was debated[S26]) and all His sidechains except for H297$^{6.52}$ neutral Hid. The calculated p$K_\mathrm{a}$ values of all residues in the apo mOR are given in our previous work.[S26] We note that H297$^{6.52}$ has a calculated p$K_\mathrm{a}$ of 6.8 in the apo mOR.

**Preparation and equilibration of the fentanyl-mOR complex.** The fentanyl-bound mOR was prepared by superimposing the docked fentanyl-mOR complex structure obtained from a previous study[S10] onto the final snapshot of the above membrane-embedded apo mOR simulation. In the resulting structure, fentanyl is bound in the orthosteric site, making a salt bridge between its positively charged piperidine amine and the negatively charged sidechain of D147$^{3.32}$. Water molecules overlapping with the ligand were removed. Since the fentanyl carries a $+1$ charge, one $Na^+$ ion was deleted to maintain the system neutrality. The fentanyl-mOR complex embedded in the membrane was energy minimized for 5,000 steps and then relaxed through a series of equilibrium simulations for a total of 63 ns. In the first 3 ns, the protein backbone and ligand heavy atoms were harmonically restrained with the force constant graduately reduced from 2.5 to 0.25 kcal/mol/Å$^2$. At the same time, the force constant on the protein sidechain heavy atoms were gradually reduced from 1.25 to 0.25 kcal/mol/Å$^2$. In the next 50 ns, all restrains were removed. The root-mean-square deviation (RMSD) of the protein backbone remained steady at $\sim$1Å, and the ligand remained bound to the orthosteric site and maintaining the salt bridge with D147$^{3.32}$. In the RMSD calculation, the trajectory frames were aligned to the starting structure based on the $C_\alpha$ atoms of the transmembrane helices (TM). The Nterminal residues 52–64, loops (residues 96-101, 131-135, 172-180, 206-224, 262-268, 306-311), and C-terminal helix (residues 337-347) connected to TM7 were excluded in the calculation.

**Well-tempered metadynamics.** We applied well-tempered metadynamics method [S11,S27] to examine fentanyl-mOR dissociation. The technique expedites ligand dissociation by depositing history-dependent biases ($V_{\mathrm{meta}}$) along a reaction coordinate,[S27] where $V_{\mathrm{meta}}$ is defined as

$$V_{\mathrm{meta}}(r(t)) = \sum_{t'=\delta t}^{t'<t} W \prod_{i=1}^{N_{cv}} exp\left(-\frac{(r_i(t) - r_i(t'))^2}{2\sigma_{r_i}^2}\right). \tag{S1}$$

Here $W$, $\sigma$, and $\delta t$ are the Gaussian height, width, and deposition time step, respectively. Free energy ($\Delta$G) can be estimated from the biases as:[S11]

$$\Delta G(r(t)) = -\frac{T + \Delta T}{\Delta T} V_{\mathrm{meta}}(r(t)) \tag{S2}$$

where $T$ is the simulation temperature (310 K) and $\Delta T$ is the tempering parameter that smoothly adjusts $W$ each time a particular point in the reaction coordinate is visited. The dissociation (residence) times of the ligand can be estimated as demonstrated by several previous studies.[S13,S14,S28,S29]

**Protocol of metadynamics simulations.** Two sets of metadynamics simulations of fentanyl-mOR dissociation were performed using the ColVars module[S12] of NAMD 2.13.[S6] In the first set of simulations, the final snapshot from the 50-ns unrestrained simulation of the docked structure was used as the starting structure, whereby fentanyl's COM $z$ is $\sim$4.5 Å with respect to the C$\alpha$ z of D147$^{3.32}$. Note, the position of this atom did not fluctuate much during the simulations. Calculated over the 15 metadynamics trajectories in the presence of Hid297, the C$\alpha$ z is 11$\pm$0.3 Å. Following the results of our recent continuous constant pH and WE simulations,[S26] neutral N$\delta$-protonated Hid297, neutral N$\epsilon$-protonated

Hie297 and doubly protonated Hip297 were examined. 48 simulations were performed for each H297 state and started with randomly assigned initial velocities. Hie297 and Hip297 tautomers were prepared by mutating residue Hid297 in the equilibrated structure using VMD.[S30] One $Na^+$ ion in the Hip297 system was removed to neutralize the system. Both systems were energy-minimized and heated to 310 K before starting the simulations. In total, 144 simulations were performed and each lasted 10-100 ns, amounting to an aggregate simulation time of $\sim$4 $\mu$s.

The first set of simulations showed two exit pathways: "direct" and "deep". For the direct pathways, the ligand escaped directly to the extracellular surface. For the deep pathways, the ligand sampled deep into the protein before exiting at a longer timescale than via the direct pathways. In a separate project, we analyzed the 40 $\mu$s weight-ensemble trajectories [S26] and found that in the most populated configuration of the fentanyl-mOR complex, fentanyl is bound to D147 with the COM $z$ $\sim$1 Å. Using this global minimum state as the starting structure, we carried out a second set of metaydynamics simulations. 15 independent runs for each of the $H297^{6.52}$ protonation states were conducted, amounting to an aggregated sampling time of $\sim$2 $\mu$s.

Fentanyl-mOR contact number (N-terminus, loops connecting TM helices, and C-terminal helix are excluded) and fentanyl's COM z with respect to the COM z of the $C_\alpha$ atoms in the orthosteric site were used as the collective variables. The orthosteric site residues were identified from the BU72-bound mOR structure (PDB: 5C1M[S7] as Y75, Q124, N127, W133, L144, D147, Y148, M151, F152, L232, K233, V236, A240, W293, I296, H297, V300, W318, H319, I322, and Y326. Following the work of Fiorin et al,[S12,S31] The contact number (CN) is defined as

$$\mathrm{CN} = \sum_{i \in \mathrm{mOR}} \sum_{j \in \mathrm{FEN}} \frac{1-(d_{ij}/4.5)^8}{1-(d_{ij}/4.5)^{16}}, \tag{S3}$$

where $d_{ij}$ is the distance between the heavy atoms $i$ and $j$ in mOR and fentanyl, respectively. The effective cutoff distance is 4.5 Å. In the metadynamics simulations, a Gaussian-based bias potential was deposited in every 10 ps, with $W$ of 0.5 kcal/mol, $\sigma$ of 0.5 Å for $z$, and 5 for the contact number. ΔT was set to 14T, following a previous study by Tiwary et al.[S13] The PLUMED program[S32,S33] was used to calculate $t_{\mathrm{real}}$ and to reconstruct the free energy surfaces projected onto the biased (i.e., z and contact number) and non-biased CVs (e.g., z, FEN–D147/FEN–H297 distance). The simulations were terminated once the ligand was completely outside of the receptor (fentanyl COM z relative to that of the orthosteric site is greater than 25 Å and contact number is nearly zero).

**Analysis and calculation of dissociation time.** To properly determine the z cutoff value for defining the unbound state, we manually examined all trajectories to avoid ligand reentry or sampling at the protein surface. Based on this and after the verification that the calculated $\tau$ and $p$ values do not significantly change beyond the cutoff (see Fig. S5), we arrived at 15 Å (fentanyl COM z relative to that of the orthosteric site) as the optimum cutoff for defining the unbound state.

After we noticed that the escape times based on all trajectories (with each protonation state of H297) cannot be fit to a single dissociation time, we found that in some trajectories (direct pathways), the ligand escaped to the extracellular surface directly from the orthosteric site region, while in other trajectories (deep pathways), it sampled in a region well below 0 Å prior to escaping the receptor. Thus, we divided the trajectories into two groups, those sampling the direct pathways (fentanyl's z never goes below -0.5 Å), and those sampling the deep pathways (fentanyl's z can go below -0.5 Å). The z cutoff was based on the examination of the approximate free energy surfaces projected onto $z$ and contact number. Later, after the completion of the second set of simulations, we found that the free energy surfaces based on the deep trajectories contain the global minimum from the weighted-ensemble simulations. By contrast, the direct trajectories did not sample the global minimum state . We also verified that the $\tau$ values are minimally affected by slightly changing the $z$ cutoff.

Following Salvalaglio et al.,[S34] the dissociation time ($\tau$), which is the reciprocal of $k_{\text{off}}$ is calculated by fitting the distributions of the unbiased dissociation times $t_{\text{real}}$ recovered from individual simulations to the theoretical cumulative distribution function (TCDF),

$$\text{TCDF} = 1 - e^{-t_{\text{real}}/\tau}. \tag{S4}$$

$t_{\text{real}}$ can be estimated using the acceleration factor ($\alpha$) defined as:[S28]

$$\alpha \; =< e^{\beta V_{\text{meta}}(r(t_{\text{bias}})} >= \; t_{\text{real}}/t_{\text{exit}}, \tag{S5}$$

where $t_{\text{exit}}$ is the biased dissociation time and $\beta$ = 1/k$_{\text{B}}$T, or by summing the time steps until reaching $t_{\text{exit}}$:[S14,S29]

$$t_{\text{real}} \; = \sum_{t_i < t_{exit}} \delta t e^{\beta V_{\text{meta}}(r(t_i),t_i)} \tag{S6}$$

where $\delta t$ is the time step for depositing the biases. The Kolmogorov-Smirnov (KS) test was used to test the null hypothesis that the sample of dissociation times extracted from metadynamics and a large sample of times randomly generated according to the theoretical probability density reflect the same underlying distribution. The $p$ value of $\geq$ 0.05 is used for accepting the TCDF.

**Table S1:** Solvent accessible surface areas of the deep-pocket residues based on the deep-insertion trajectories as compared to the apo simulations

| Residue | z (Å) | Deep insertion state (Å$^2$) | Apo (Å$^2$) |
|---|---|---|---|
| I155 | $\sim$-9 | 25.0$\pm$8.3 | 14.1$\pm$4.7 |
| L158 | $\sim$-14 | 26.7$\pm$10.8 | 15.4$\pm$4.7 |
| P244 | $\sim$-8 | 11.9$\pm$5.4 | 3.5$\pm$2.7 |
| F289 | $\sim$-10 | 42.7$\pm$16.3 | 15.9$\pm$6.2 |

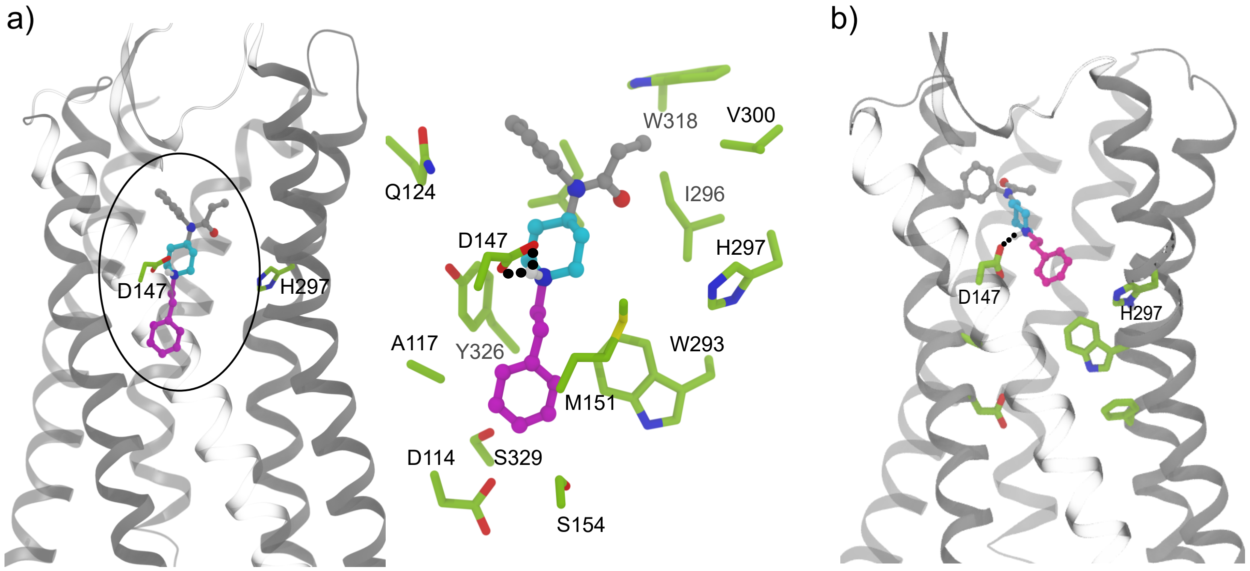


**Figure S1: Starting structures for the two sets of metadynamics simulations.** a) The most populated configuration of the fentanyl-mOR complex taken from our recent 40-$\mu$s WE simulations.[S26] Fentanyl is positioned at z∼1 Å in this structure. b) The "relaxed docked structure". Fentanyl is positioned at z∼4.5 Å in this structure. Note, the origin of $z$ is placed on the C$\alpha$ atom of D147, and both starting structures contain a salt bridge between fentanyl's piperidine nitrogen and D147's carboxylate. The phenylpropanamide, piperidine, and phenethyl groups are colored gray, cyan, and magenta, respectively. The displayed residues in the zoomed-in view are within 5 Å of fentanyl.

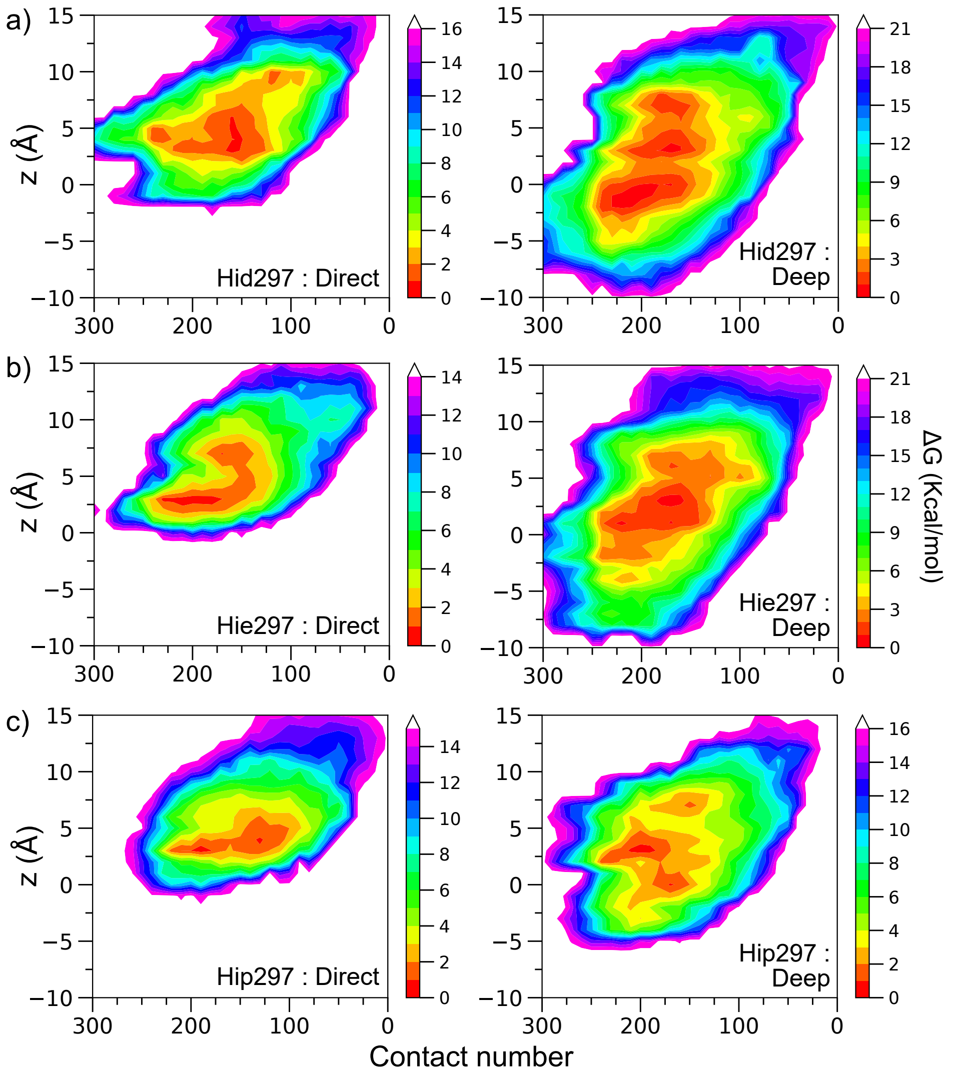


**Figure S2: Approximate free energy surfaces projected onto fentanyl's contact number and z position obtained from the simulations initiated from the relaxed docked state.** Left panel: FE surfaces calculated from the trajectories with Hid297 (**a**), Hie297 (**b**), and Hip297 (**c**), showing the dissociation of fentanyl directly from the orthosteric site. Right panel: FE surfaces calculated from the trajectories with Hid297 (**a**), Hie297 (**b**), and Hip297 (**c**), showing that fentanyl sampled well below the orthosteric site. Each of the displayed FE surface was calculated from the Boltzmann average of the FE surfaces of individual trajectories.

The free energy surfaces based on the individual trajectories were calculated using a reweighting protocol[S35] in PLUMED.[S33] A trajectory is defined as following a "deep pocket pathway" if z decreases to below -0.5 Å at any point during the simulation and "direct exit pathway" if otherwise.

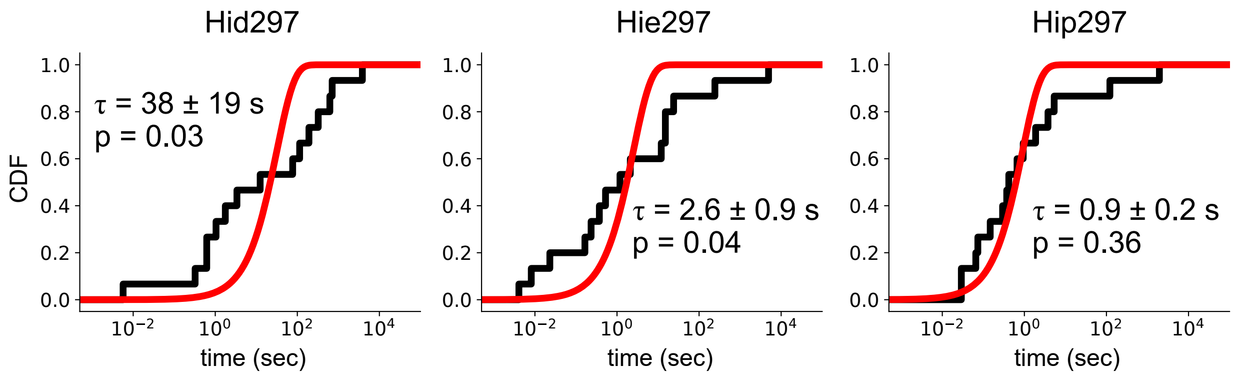


**Figure S3: Poisson fitting analysis of fentanyl-mOR dissociation time based on the simulations initiated from the global minimum state.** Cumulative distribution function (CDFs) obtained from the 15 trajectories in the presence of Hid297 (left), Hie297 (middle), or Hip297 (right). Red curve is the best fit to the theoretical CDF (Eq. S4). The calculated residence time ($\tau$) and $p$-value from the two-sample Kolmogorov-Smirnov (KS) test are given. The values of $\tau$ and standard deviations ($\pm$) were calculated from the bootstrapping analysis for $n$ times over $n-1$ samples, where n is 15.

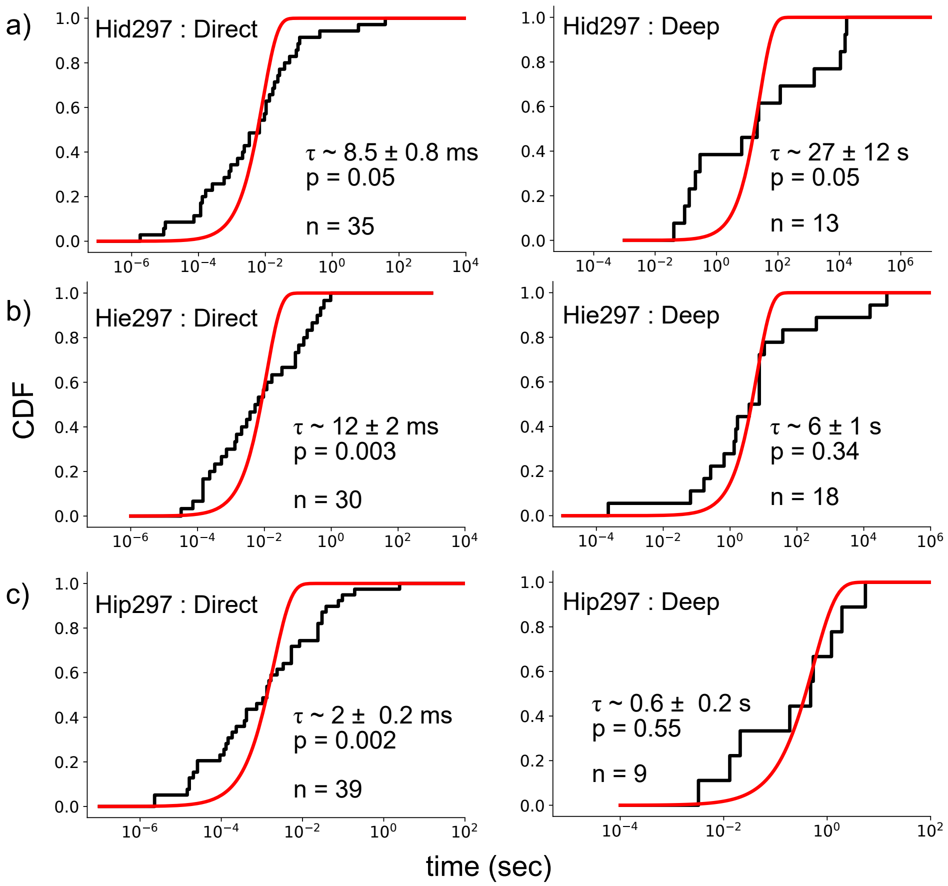


**Figure S4: Poisson fitting analysis of the fentanyl-mOR dissociation times based on the simulations started from the docked structure.** Cumulative distribution functions (CDFs) obtained from the trajectories in the presence of Hid297 (a), Hie297 (b), or Hip297 (c). Trajectories in which fentanyl escapes directly from the orthosteric site region are shown on the left and those that can sample a global minimum are shown on the right.

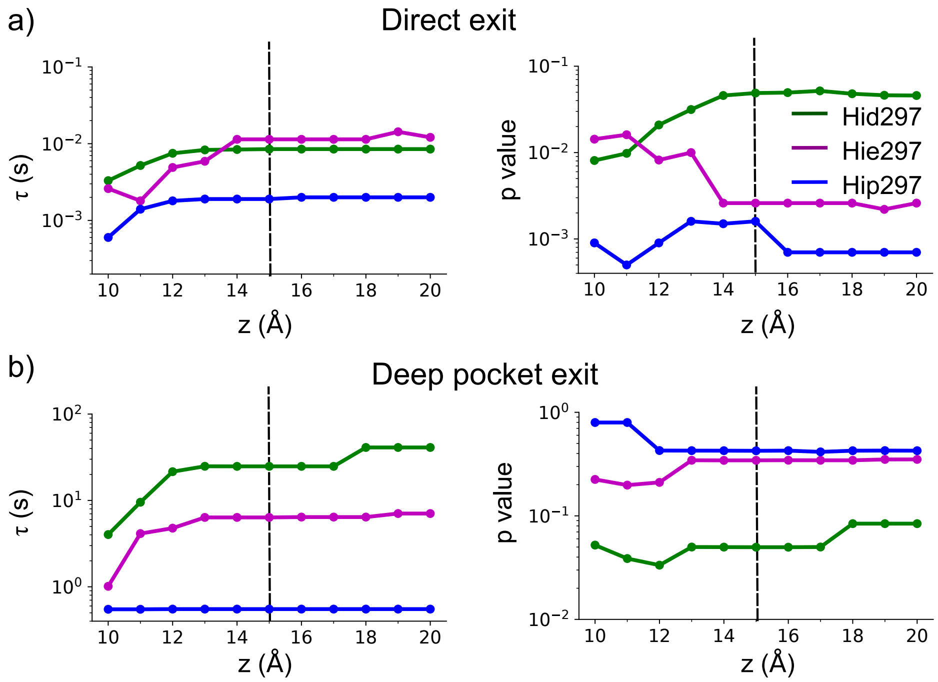


**Figure S5: Calculated dissociation time and $p$ value versus the $z$-position cutoff used to define the unbound state.** Simulations started from the relaxed docked structure were used. a) Data from the direct exit trajectories. b) Data from the deep pocket trajectories. The unbound state was defined as $z$ above 15 Å (dashed line), where fentanyl reaches the extracellular surface. Note, in most cases, $\tau$ and $p$ values do not change significantly beyond this cutoff.

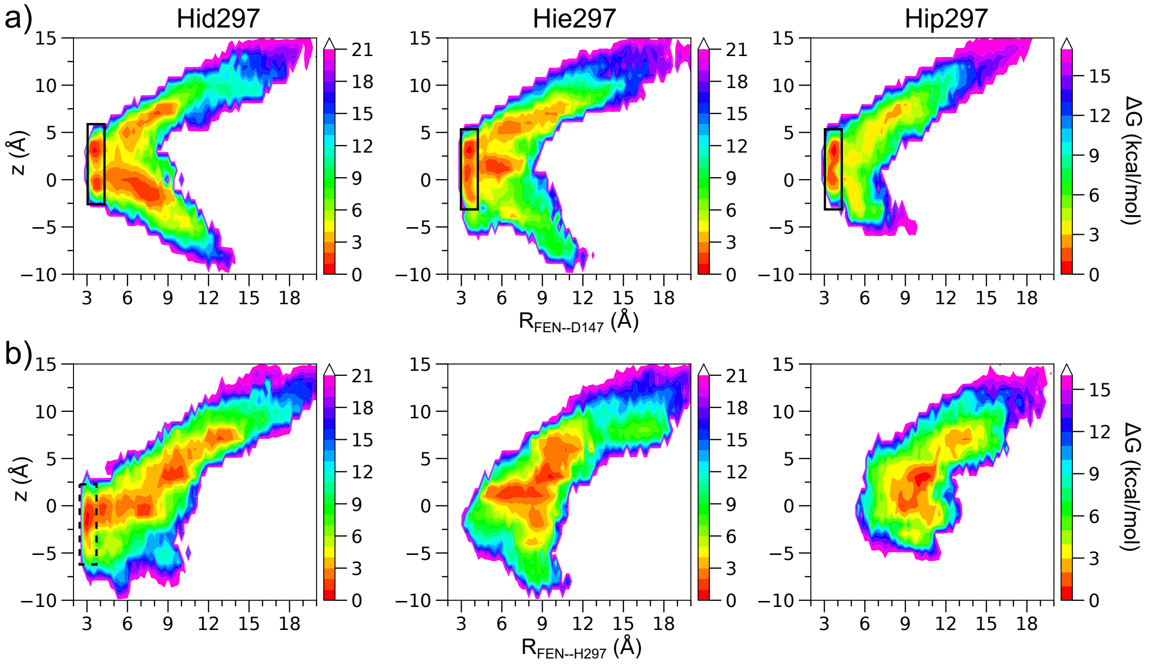


**Figure S6: Free energies as a function of fentanyl's z position and distance to D147 or H297 based on the simulations initiated from the relaxed docked structure.** a) Free energy as a function of fentanyl's z and the distance between piperidine:N and D147:C$\gamma$ based on the Hid, Hie, or Hip trajectories. b) Free energy as a function of fentanyl's z and the minimum distance between piperidine:N and H297:N$\delta$/N$\epsilon$ based on the Hid, Hie, or Hip trajectories. The Boltzmann average of the free energy surfaces of the individual deep pocket trajectories was taken. The latter were calculated using a reweighting protocol[S35] in PLUMED.[S33] The origin of $z$ is placed on the C$\alpha$ atom of D147. The black box highlights the region where the piperidine–D147 salt bridge is formed. The dashed box highlights the region where the piperidine–H297 hydrogen bond is formed.

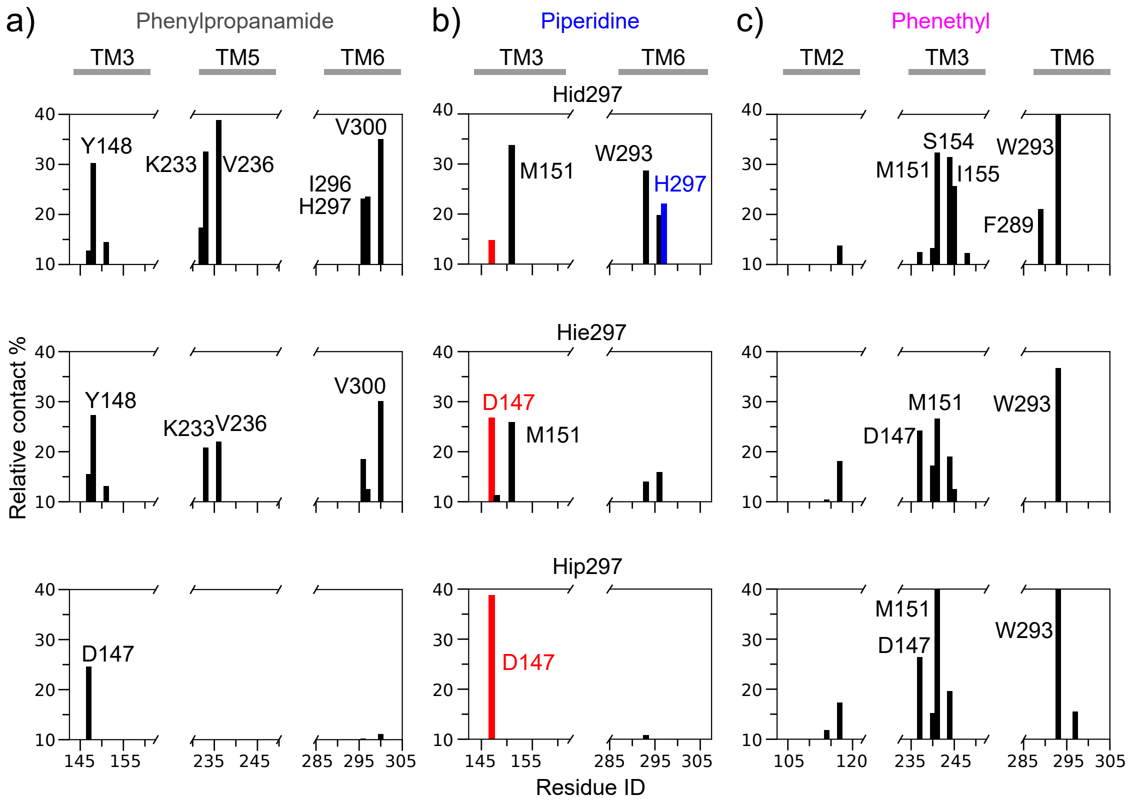


**Figure S7: Fentanyl-mOR contact profile from the simulations initiated from the relaxed docked structure. a,b,c)** Relative fractions of contacts between mOR residues and the phenylpropanamide (**a**), piperidine (**b**), or phenethyl (**c**) group calculated from the deep vs. direct trajectories in the presence of Hid297 (top), Hie297 (middle), or Hip297 (bottom). Only positive relative contact fractions $\geq$10% are displayed. Residues with relative contact fractions $\geq$20% are labeled. The contacts are defined using a 4.5-Å distance cutoff between heavy atoms.
